# A central component of the N1 event-related brain potential could index the early and automatic inhibition of the actions systematically activated by objects

**DOI:** 10.1101/341057

**Authors:** Molly Touzel, Christine Snidal, Julia Segal, Louis Renoult, J. Bruno Debruille

**Author notes:** Corresponding author, Douglas Mental Health University Institute, 6875 Blvd LaSalle, Montréal, (Qc) H4H 1R3, Canada.

## Abstract

Stimuli of the environment, like objects, systematically activate the actions they are associated to. These activations occur extremely fast. Nevertheless, behavioural data reveal that, in most cases, these activations are then automatically inhibited, around 100 ms after the occurrence of the stimulus. We thus tested whether this early inhibition could be indexed by a central component of the N1 event-related brain potential (ERP). To achieve that goal, we looked at whether this ERP component is greater in tasks that could increase the inhibition and in trials where reaction times happen to be long. The illumination of a real space bar of a keyboard out of the dark was used as a stimulus. To maximize the modulation of the inhibition, the task participants had to performed was manipulated across blocks. A look-only task and a count task were used to increase inhibition and an immediate press task was used to decrease it. ERPs of the two block-conditions where presses had to be prevented and where the largest central N1s were predicted were compared to those elicited in the press task, differentiating the ERPs to the third of the trials where presses were the slowest from the ERPs to the third of the trials with the fastest presses. Despite larger negativities due to motor potentials and despite greater attention likely in immediate press-trials, central N1s were found to be minimal for the fastest presses, intermediate for the slowest ones and maximal for the two no-press conditions. These results thus provide a strong support for the idea that the central N1 indexes an early and short lasting automatic inhibition of the actions systematically activated by objects. They also confirm that the strength of this automatic inhibition spontaneously fluctuates across trials and tasks. On the other hand, just before N1s, parietal P1s were found greater for fastest presses. They might thus index the initial activation of these actions. Finally, consistent with the idea that N300s index *late* inhibition processes, that occur preferentially when the task requires them, these ERPs were quasi absent for fast presses trials and much larger in the three other conditions.

**Highlights:** Event-related brain potentials (ERPs) elicited by a real object

Smaller parietal P1s and greater central N1s for slowest than for fastest motor responses

Even greater central N1s for tasks without such responses

Central N1s may index early inhibition of stimulus-activated actions

N300s could index late inhibition of stimulus-activated actions

## Introduction

Quite often now, affordances is a word used to designate the possible *actions* that are systematically and pre-consciously primed by objects (Tucker and Ellis, 1998)^1^. Such actions are now central to theories of embodied cognition (for a recent review, see Matheson, White & McMullen, 2015). Systematically activated ‘motor representations’ of objects are considered integral to the identification of an object (Clark, 1999; Wilson, 2002). These systematic activations were shown by Tucker and Ellis (1998), as well as others (e.g., Borghi & Riggio, 2008; Phillips & Ward, 2002), in experiments in which subjects viewed *pictures* of graspable objects whose handle was oriented either to the left or to the right. Responses were faster with the left hand for pictures of objects with handles oriented to the left whereas right-hand responses were faster when the handle of the objects was oriented to the right. Although these compatibility effects seem to exist even when attention is not focused on affordances (Symes, Ellis & Tucker, 2006), their sizes are increased by attending to the action-related features of the object (Tipper, 2010). For instance, focusing on a cup’s handle, elicited larger compatibility effects than focusing on unrelated features, such as the color of the object (Tipper, Paul & Hayes, 2006).

Imaging studies corroborated the idea that objects systematically activate sensory-motor brain regions and thus motor representations. A PET study of object’s perception showed activations in the inferior parietal lobule, the supplementary motor area, the dorsal and ventral precentral gyri and the cerebellum in comparison to viewing non-objects (Grèzes & Decety, 2002). Studies using manipulable objects have shown that the observation of tools, even without having to use them, trigger strong left dorsal premotor cortex activation (Grafton, Fadiga, Arbib & Rizzolatti, 1997). The ventral premotor cortex and the left posterior parietal cortex were also activated by viewing and naming pictures of tools compared to control items in a study by Chao and Martin, (2000). Other works have provided evidence for a functional separation of the dorsal pathway (Rizzolatti, Luppino, & Matelli, 1998). It has been divided into a dorso-dorsal subpathhway that controls actions and a ventro-dorsal pathway that codes the representations of actions (Binkofski & Buxbaum, 2013; Buxbaum & Kalenine, 2010; Rizzolatti & Matelli, 2003). The former runs from visual areas (V6) and the superior parietal lobe to the dorsal premotor cortex. The latter projects from the medial superior temporal area into the inferior parietal lobe.

However, most of the times, performing the actions that are systematically primed by the occurrence of stimuli is contextually inappropriate. This is true not only for trying to grab an object whose picture appears on a screen but also, in everyday life, for acting in a way that would not be consistent with our goals. Given that all objects of our environment seem to prime the actions they are associated to, inhibition of all those unwanted actions must be a quasi ceaseless mechanism of wide importance (Miller, 2000; Ridderinkhof, 2002a, 2002b; Ridderinkhof, van den Wildenberg, Wijnen, & Burle, 2004; Sumner et al., 2007). Many facts are consistent with this view. For instance, when subjects have to press some buttons on a board while ignoring others, the motor representations associated with the to-be-ignored buttons are inhibited (e.g., Tipper, Lortie & Baylis 1992). Or, when certain lesions disrupt the inhibition of affordances, in a study of anarchic limb behaviors for example, the patient would act upon the affordance to grasp a cup with the hand congruent with handle orientation, rather than following the instruction to always grasp with the hand closest to the cup (Riddoch, Edwards, Humphreys, West & Heafield, 1998). Similarly, frontal lesions result in ‘utilization behaviors’, which are defined as the difficulty in resisting the use of an object within reach (e.g., Archibald, Mateer & Kerns, 2001). When discovering the pen of the doctor on his/her desk, these patients will grab it and use it.

The results obtained in yet another tradition of research objectivize the existence of such a systematic inhibition of actions. These results show that selective motor inhibition occurs automatically even without conscious perception and voluntary effort (Eimer & Schlaghecken, 2003; Sumner et al., 2007; Atas & Cleeremans, 2015). They were obtained by using brief presentations of elementary visual objects (primes), which were immediately masked by another visual stimulus before a target appears. The primes were either compatible or incompatible with the target. In this type of design, *compatible* primes *delayed* reaction times (RTs) to targets. Recent works show that these RT increases reveal an automatic motor inhibition rather than a competition with an incorrect “opposite-to-prime” response (Vanio et al., 2011; Ocampo and Finkbeiner, 2013; Atas & Cleeremans, 2015).

Electrophysiological studies of object affordances have shed light on the mechanisms underlying the inhibition of affordances. The disappearance of a graspable object was found to enhance mu-rhythm synchronization (Schuck et al., 2010) while its viewing had induced a mu-rhythm *de*synchronization. In a quite different domain, a study by Debruille et al. (2012) investigated the inhibition of the social affordances that are automatically elicited by faces, such as making eye contact, saying hello and starting a conversation. For that purpose, they used the face of a confederate with whom making eye contacts was appropriate and the face of a dummy for whom this automatically activated affordance had to be inhibited. A block design was used in order to increase the focus on activations that occur systematically and would thus have to be automatically inhibited when inappropriate. Event-related brain potentials (ERPs) elicited by the two face-stimuli were then compared. Those of the dummy included larger central N300s than those of the face of the confederate. This was interpreted as indexing the greater amount of affordance inhibition required for the dummy. Interestingly, these ERPs had the same latency and the same central scalp distribution as the NoGo N2 potential (Falkenstein et al., 1999; Ocklenburg et al., 2011), an ERP that is precisely elicited in conditions where (simple) actions associated to simple stimuli have to be prevented and which has thus been associated to their inhibition (Bruin and Wijers, 2002; Jodo and Kayama, 1992; Roche et al., 2005).

Nevertheless, these N300- and No-Go-N2-ERPs onset around 200 ms post stimulus onset. They thus start too late to index mechanisms that could prevent the actions systematically activated by the stimulus. Indeed, these systematic activations occur extremely early. For instance, when participants are observing graspable objects, the increase in muscle excitability starts as early as 120 ms after the onset of the presentation (Franca et al., 2012). Similar delays were obtained in a study where participants learned to associate the contraction of the tibialis anterior muscle of each ankle in responses to an auditory cue (Schneider et al., 2004) or when measuring facial action (i.e., spontaneous mimicry) in response to the presentation of happy and neutral faces (Korb, Grangean and Sherer, 2010). Actions associated to a stimulus thus appear to be activated in the brain in as little as 100 ms after the onset of the stimulus. Indeed, at least two dozens of milliseconds have to be subtracted from the 120 ms delays mentioned above in order to take into account the time, for the brain output, to reach and excite the muscles recorded and, for the first two studies, for the movement to attain a measurable size. Brain mechanisms that prevent affordance from being acted upon thus have to start as early as 100ms. Without such mechanisms, we could start acting each time a stimulus associated to an action occurs in our environment. This can also be deduced from the duration of the presentation of the primes in the studies mentioned above to show that selective motor inhibition occurs automatically even without conscious perception and voluntary effort (Eimer & Schlaghecken, 2003; Sumner et al., 2007; Vanio et al., 2011; Ocampo and Finkbeiner, 2013; Atas & Cleeremans, 2015). The elementary visual objects used as primes were presented for only 30 to 70 ms before being masked.

The high temporal resolution of the event-related brain potential (ERP) technique makes it an ideal tool for exploring such a fast inhibition mechanism. The N1 ERP elicited by visual stimuli onsets at a latency compatible with the early inhibition necessary to prevent immediate action upon fast affordance activation. Although its posterior, that is, occipital components, are known to index processing of the physical features of the visual stimuli, the functional significance of its central components is not well known. Most interestingly, these more anterior components have been previously related to the processing of affordances in two studies. The first, that of Proverbio, Del Zotto and Zani, (2007), reports that manipulable objects elicited larger central N1s than animal entities, suggesting that anterior motor regions might be involved in action-centered representations of tools. The second is that of Debruille et al. (2012) whose N300 results have been mentioned above. In this study, ERPs in the N1 time windows were more negative for the dummy, with whom social affordance were inappropriate, than for the real person. Also noteworthy, the N1 has been shown to be of greater amplitude for stop-than for go-signals (Raud & Huster, 2017; Senderecka, 2016; Kenemans, 2015) and for successful than for unsuccessful stop-signal conditions (e.g., Bekker et al. 2005; Wild-Wall et al. 2008; Pires et al. 2014). It has thus been proposed as an index of a *directed* (or learned) action-inhibition.

The first goal of the present study was thus to test whether the central N1 could index the early and *automatic* inhibition that objects would spontaneously trigger just after they activate the actions they are associated to. This early automatic inhibition was hypothesized to be of very short duration and to last only until the occurrence of the second negativity, that is, the N300 or the No-Go N2, which seems to index a later, more context-dependent, inhibition.

The second goal of the current study was to search for a candidate ERP index of the fast and systematic activation of actions (or affordances) that occurs before the automatic inhibition. We thus explored whether the ERP that precedes it, that is, the P1, could index it. This idea was based on the parieto-central P1s that were related to initial action activation by Kiefer et al. (2011). Here, we assumed that greater activations would lead to faster actions. We thus hypothesized greater parietal P1s for trials with short than for those with long reaction times, as the threshold for real action would be reached faster.

The third goal was to examine the N300 elicited by the display of an object and see whether its amplitude would vary as an index of the late and context dependent inhibition of the actions automatically activated by stimuli, as seemed to be the case for the face of the dummy in Debruille et al. (2012). Accordingly, we looked at whether this amplitude would be large in the tasks where subjects have to prevent performing the action than in tasks where they have to perform it.

To achieve these three goals, we designed a study with one simple and highly familiar object: the spacebar of a computer keyboard. It was assumed that, for anyone who grew up in the age of computers, this stimulus systematically activates the action of pressing it, just as a coffee cup handle affords grasping (an idea already used in other recent studies, e.g., Horváth, 2013). The present study was conducted in a completely dark room and the spacebar of a keyboard placed in front of the subject was illuminated sporadically, thus appearing out of darkness. The method used to manipulate the automatic inhibition was based on the fact that some automatic mechanisms (e.g., those of the knee reflex) can be influenced and the action, prevented, reduced or, on the contrary, amplified. A block design was used to maximize this modulation. The space bar was the only stimulus used and participants had to process it in exactly the same way throughout each block to reinforce their strategy. Two tasks were used to maximize the inhibition of presses, a look only task and a count the number of appearance task. A task where participant had to immediately press the space bar upon its appearance was used to decrease automatic inhibition. Larger central N1s were thus predicted in the two former tasks than in this latter one.

Given that participants’ attention spontaneously fluctuates across the trials of an experiment, it was assumed that both the strength of the systematic activation and the strength of the early automatic inhibition would also fluctuate across trials. In the immediate press task, these fluctuations should lead to faster presses when activation is strong and inhibition weak and to slower presses when activation is weak and inhibition strong (somewhat like in Atas & Cleeremans, 2015). The ERPs elicited by the illumination of the space bar were thus averaged separately according to whether the press was slow or fast. Larger central N1s were predicted for the former than for the latter.

Importantly, two very well-known effects should drive N1 amplitudes in the direction opposite to these predictions. Firstly, the attention effect, as, relative to the look-only and the count task, the immediate press task stimulates vigilance as participants have to press as fast as possible. It should thus be associated to greater attention and thus to greater N1s, an ERP effect known to be maximal at occipital electrode sites (Hillyard, & Anllo-Vento, 1998; Johannes et al. 1995; Hillyard et al. 1995; Luck, Fan & Hillyard, 1993). Secondly, the motor potentials elicited by press trials. They will make ERPs more negative at central sites over the hemisphere contralateral to the pressing-hand just before the pressing (Coles, Gratton & Donchin, 1988; Miller & Hackley 1992). This could thus be mistaken for greater central N1s as reaction times in simple tasks where no choice has to be made are known to be roughly between 200 and 300 ms (Niemi & Näätänen, 1981). The readiness potentials that precede them will thus be within the N1 time window. In these conditions, observing greater central N1s for the two tasks where the pressing had to be prevented would thus be all the more demonstrative for the inhibition hypothesis.

On the other hand, in order to look for an index of the prior activation, we also compared ERPs of the four conditions (i.e., fast-presses, slow-presses, count and look only) in the time window preceding these N1s. Finally, N300s were explored to see if their amplitudes would also be minimal when subjects act upon the activated affordance, that is, in the press task.

## Methods

### 1. Participants

Nine females and 11 males with a mean age of 24 (range 18 to 30) were recruited as participants. They all learned about the experiment through classified ad websites. Right-handed, they had at least some university education and had normal- or glasses-corrected to normal vision. None reported any personal or family history of severe neurological or psychiatric disorder. Participants were excluded if they consumed more than twelve drinks of alcoholic beverages per week or if they used recreational drugs, except if they used marijuana less than once per week. They read and signed an informed consent form approved by the Douglas Institute Research and Ethics Board and were compensated for their time $15 per hour.

### 2. Procedure

Subjects performed the tasks seated in front of a computer in a completely dark room. The screen and the keyboard of the computer were concealed with cardboard, such that when the screen of the computer turned from black to white, its emanated light, which lasted 500 ms, was directed onto the spacebar so that only this object (and the cardboard) was visible. Inter-stimulus intervals varied randomly between 1.4 and 2.3 seconds. During these intervals, the screen of the monitor was black and subjects could not see the spacebar or any other object, given the total darkness of the room. All 20 subjects completed an experiment that consisted of 3 block-tasks, each one including 100 occurrences of the space bar. They were instructed to rest their right index finger just in front of the spacebar so as to be ready to press it without obscuring its illumination. In the ‘Press’ task, subjects pressed the spacebar as fast as possible as soon as it appeared. In the ‘Count’ task, they gazed at the spacebar and counted the number of times it appeared, giving a final report of their count at the end of the block. In the ‘Look only’ task, subjects looked at the spacebar as it appeared and disappeared and were instructed to never direct their gaze away from it. The order of these three block-tasks was counterbalanced across subjects.

### 3. Data Acquisition

The electroencephalogram (EEG) was recorded with tin electrodes mounted in an elastic cap (Electrocap International) at 28 of the sites of the extended International 10-20 system (Electrode Position Nomenclature Committee, 1991). The reference electrode was placed on the right earlobe. The active electrode sites were grouped in a sagittal subset, which included Fz, FCz, Cz and Pz; a parasagittal subset, which comprised FP1/2, F3/4, FC3/4, C3/4, CP3/4, P3/4, and O1/2; and a lateral subset, which consisted of F7/8, FT7/8, T3/4, TP7/8 and T5/6. The EEG was amplified 20,000 times by Contact Precision amplifiers. The half amplitudes cut-offs of their high- and low-pass filters were set at 0.1 and 100 Hz, respectively, with an additional electronic notch filter to remove 60 Hz contamination. Signals were digitized on-line at a 256 Hz sampling rate and stored for subsequent averaging using the I-wave (version 5.24) software package. Behavioral data, consisting of response times for the press-task, were recorded using the E-prime (version 2.0) software.

### 4. Data Processing

EEG epochs of the 100 trials of each block-task were examined. Trials of the press-task in which response time exceeded 500ms or in which there was no response were excluded, leading to an average of 5.6 trials being excluded. Trials contaminated by eye movements, excessive myogram, amplifier saturations or analog to digital clipping were removed off-line when analog to digital clipping exceeded a 100 ms duration or when the absolute value of the amplitude exceeded 100µV. For the press-task, the third of the trials in which subjects responded the fastest were separated from the third of the trials in which subjects responded the slowest. ERPs to stimuli were computed in each subject in the ‘Count,’ ‘Look Only,’ ‘Fast Press’ and ‘Slow Press’ conditions by averaging the EEG epochs of these trials in each task using a -200 to 0 ms baseline and looking up to 600 ms post stimulus onset.

### 5. Measures and Analyses

Visual inspection of event-related potentials (ERPs) revealed differences between conditions in the P1, N1 and N300 time windows. To assess the statistical significance of these differences, mean ERP voltages were first measured in time windows centered on their peaks, that is, in the 80-120, the 120 to 200 and the 300 to 500 ms time window, respectively. These measures were entered in three repeated-measures ANOVAs using a multivariate approach. The ANOVAs for the sagittal subset of electrodes included two within-subject factors a) condition, which had 4 levels: ‘Fast Press’ vs. ‘Slow Press’ vs. ‘Count’ vs. ‘Look Only’) and b) electrode site. In the ANOVAs dedicated to the parasagittal and lateral subset of electrodes, a hemiscalp (left vs. right) factor was added. Other ANOVAs were then conducted on the region of interest (e.g., C3, Cz & C4 for the central N1s). To test the hypothesis of a difference between fast and slow presses, an additional ANOVA was conducted comparing only the fast-press and the slow-press conditions. A similar ANOVA was also conducted to determine if there was any difference between ‘Count’ and ‘Look Only’ conditions. The Greenhouse and Geisser (1959) correction for lack of sphericity was used to correct degrees of freedom for the factors that had more than two levels (i. e., condition & electrode). In this case, original degrees of freedom are reported with corrected p-values.

## Results

### 1. Behavioral Data

Average response times in the press condition were 206 ms (SD=35.9) for the 3^rd^ of the trials which induced the fastest responses and 291 ms (SD=34.5) for the 3^rd^ of the trials which induced the slowest ones. In the ‘Count’ task, subjects reported an average of 100.5 illuminations (SD=2.6) out of the 100 illuminations of the space bar.

### 2. Electrophysiological data

#### 2.1. Visual inspection

Given that block design experiments are now much less frequently used, ERPs will first be described qualitatively in order to pinpoint their particular characteristics. Figure 1, where negative polarity is plotted upward, shows the unsmoothed grand average ERPs of the four conditions. It first displays an early difference (e.g., at Pz & P4) where parietal P1s seem larger for fast presses (green lines) than for other conditions. Then, in the following time window, at sagittal and right para-sagittal sites, the first sizeable negative deflections, the N1s appear smaller for these fast presses, intermediate for slow presses (red lines) and maximal for the count- (blue lines) and the look-only- (black lines) block-condition. Importantly, these smallest N1s for fast presses appear unlikely to be due to larger P1s, as the latter were maximal at parietal sites. At more anterior sites (e.g., C4, FC4 & FCz), no larger P1s can be seen whereas N1s for fast presses clearly appear smaller than N1s for other conditions. This contrasts with what can be observed at occipital sites, where N1s look larger for both press-trials than for the two other conditions, especially at O1. These N1s are followed by a P2, which can be best individualized at Fz and FCz, around 200 ms. Later, peaking around 300 ms, is a large positive wave maximal at Pz, which clearly appears to be a P3b occurring before the second negative deflection, as can be seen in simple tasks where only one stimulus occurs repeatedly and where no choice has to be made (Donchin et al. 1978). This late posterior positivity appeared much larger for fast presses than for the three other conditions, in line with the well-known sensitivity of this potential to task relevance and attention. These P3bs precede sizable negative waves (second negativities) maximal around 400 ms at Fz, which seemed almost abolished for fast-presses and large for all other conditions. These latter differences appeared maximal at FCz, Cz and were smaller at Pz.

**Figure 1.**
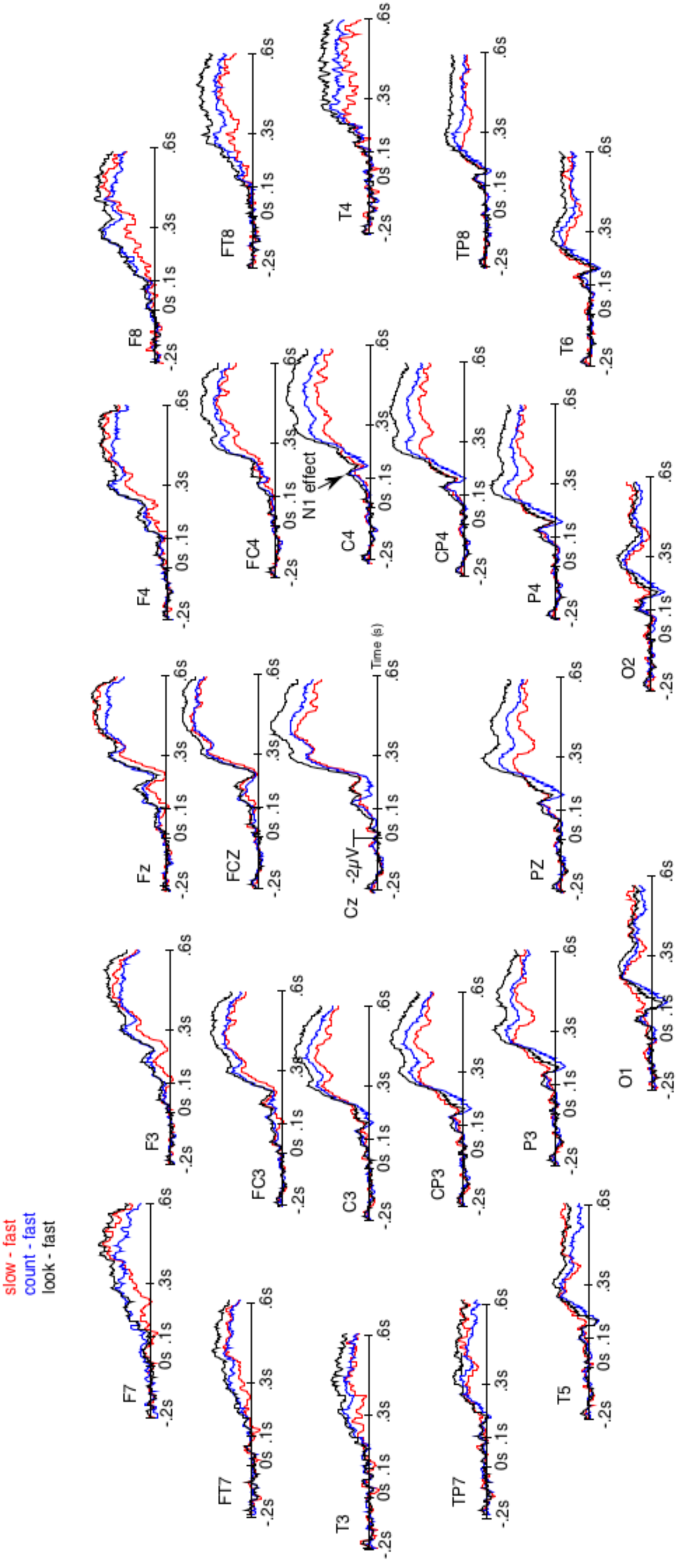
Unsmoothed grand average (n=20) of the event-related brain potentials elicited by the occurrence of the space bar of a keyboard out of the dark. Negative polarity is plotted upward. Green lines are for the fastest reaction-times trials of the block-task where the subjects had to press the space bar as fast as possible. Red lines are for the slowest reaction-times trials of that task. Blue lines are for the block-task where participants had to count mentally the number of appearances of that space bar. Black lines are for the block-task where they just had to look at the space bar. The order of these three tasks was counterbalanced across subjects.

Given that the larger P3b for fastest presses could occur earlier in some participants than in others, they could overlap N1s. There was thus a possibility for the N1s amplitude to appear small for fast presses not because of weaker N1 processes. To eliminate this possibility, we subtracted the ERPs of the fast presses from the ERPs to every other conditions. Figure 2 illustrates the results of these subtractions. There, the effect on the N1s correspond to an early deflection that clearly differs form the one corresponding to the large effect on latter ERPs. Moreover, at some electrodes sites (i.e., FCz, C4, CP4 & P4) these two deflections are separated by a return to the baseline.

**Figure 2.**
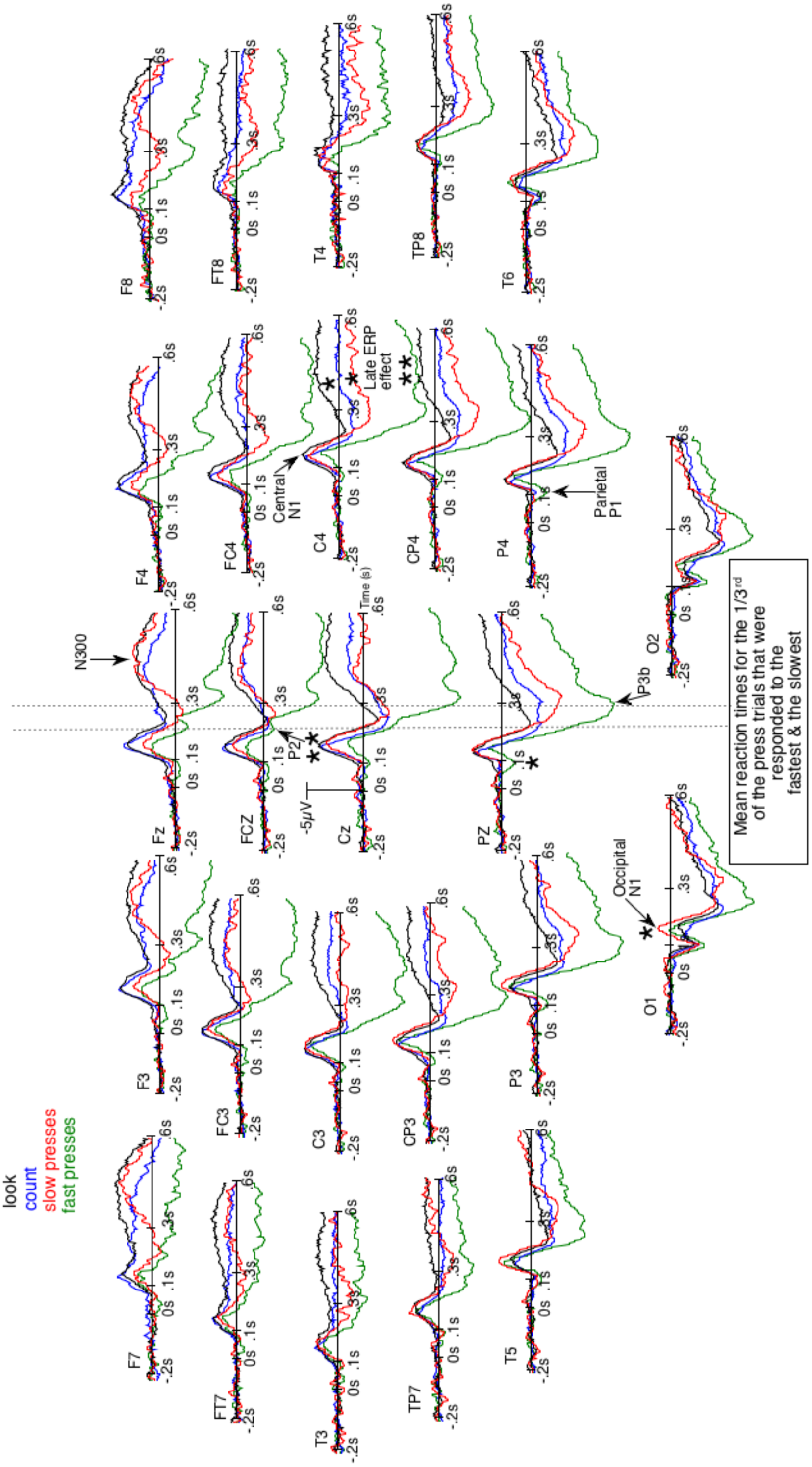
Results of the unsmoothed subtractions of the grand averages of the event-related brain potentials (ERPs) elicited by the occurrence of the space bar when reaction-times (RTs) were the fastest from the ERPs when RTs were the slowest (red lines), from the ERPs in the count block-task (blue lines) and from the ERPs of the look-only block-task (black lines). These subtractions were performed to distinguish the N1-effect from the effect on late ERPs.

#### 2.2. Analyses

##### Parietal P1 time window (80-120ms)

The ANOVA conducted with the mean ERP voltages at in the 80-120 ms time window for the sagittal subset of electrodes did not reveal any significant main effect of condition nor any significant interaction of this factor with electrode. Given that the *a priori* hypothesis was for the parietal site, a one-way ANOVA was then attempted at Pz. It founds that the P1s tended to be greater for fast press trials than for the tree other conditions at Pz (F(3,57) = 2.9, p=0.063). Similarly, the ANOVA conducted for the parasagittal subset did not reveal any significant main effect of condition nor any significant interaction of this factor with electrode or hemiscalp. Given the hypothesis and the visual inspection, a one-way ANOVA was then conducted at P4. It found that P1s were marginally greater for fast press trials than for the tree other conditions (F(3,57) = 3.0, p=0.044). The ANOVA performed for the lateral subset did not reveal any statistically significant difference.

##### N1 time windows (120-200 ms)

The ANOVAs revealed a main effect of condition at the sagittal (F(3,57)=6.8, p=0.0005), parasagittal (F(3,57=6.6, p=0.0007) and lateral (F(3,57)=7.0, p=0.0004) subsets of electrodes. Significant interactions between condition and electrode were observed only for the parasagittal subset (F(18,342)=2.9, p=0.03) where differences appeared somewhat reversed at occipital sites (O1/2). Analyses run at the sites of interest for the central N1 (FC3/4, C3/4 & CP3/4) revealed similar results for the main effect of condition (F(3,57=5.3 p=0.003), which interacted with hemiscalp (F(3,57=3.2, p=0.045) in accordance with the much smaller differences between condition seen over the left- than over the right-hemiscalp on Figure 1. The post hoc analyses run for each hemiscalp at these electrode sites to find the source of this condition x hemiscalp interaction revealed a significant effect of condition over the right hemiscalp (F(3,57=7.5, p=0.00025) and a barely significant one over the left hemiscalp (F(3,57=2.9, p=0.04). Further analyses showed greater central N1s at these sites (Fc4, C4 & CP4) for slow- than for fast-press-trials (F(1,19)=8.5, p=0.009), greater central N1s for look-only N1s than for the fast-press condition (F(1,19)=11.4, p=0.003. Conversely, ANOVAs including just the ‘Count’ and ‘Look Only’ conditions revealed no significant effect of condition and thus, no significant difference in N1 amplitude between these two tasks, at any subsets.

Another post hoc ANOVA was conducted in the same 120-200 ms time window at occipital electrodes (O1/2) to test the reversion and thus to see whether posterior N1s were greater for the task requiring more attention, that is, for the press task. This was somewhat the case (F(3,57)=3.21, p=0.05). This modestly significant effect was slightly greater over the left than over the right hemiscalp (F(3,57)=3.22 p=0.046).

##### N300 time windows (300-500 ms)

The ANOVAs revealed significantly greater N300s at the sagittal (F(3,57)=16.9, p<0.00001), parasagittal (F(3,57)=14.8, p<0.00001) and lateral (F(3,57)=10.8, p=0.00001) subset of electrodes for the three conditions other than that of the fast presses. This effect tended to interact with electrode F(18,342)=2.9, p=0.06 and with electrode and hemiscalp F(18,342)=2.4, p=0.022). No post hoc was run because of the large size of the expected impact of the late posterior positivity (LPP or P3b, see Figure 1 at Pz) during this time window. This impact makes the interpretation of the results difficult as both effects would go in the same direction. In effect, smaller N300s (or N2b) and larger P3b were predicted in the case of the press task relative to the look-only condition.

## Discussion

To test whether the amplitude of the central N1 event-related brain potential (ERP) indexes the strength of the early and automatic inhibition of the actions systematically activated by real objects, we used a novel paradigm in which the space bar of a computer keyboard was illuminated in order to appear out of the dark and to activate the action of pressing it. A block task-design was used to maximize the modulation of automatic inhibition. In each block, participants had to process the only stimulus, the space bar, in exactly the same way to re-enforce the processing strategy. Two of the three blocks were aimed at increasing inhibition. There, the task of subjects was to prevent themselves from pressing the space bar. In one of these two blocks, they had to count the number of times it appeared. In the other, they just had to look at it. The third block was used to decrease the automatic inhibition of the action. There, the task was to press the space bar as fast as possible. On the other hand, just as attention, the strength of activations and inhibitions was assumed to fluctuate from trial to trial within the same block and presses were assumed to be performed faster when their activations were stronger and their inhibitions weaker. The third of the trials where participants pressed it the fastest were thus isolated from the third of the trials where they pressed it the slower in order to see if central N1 amplitudes would be minimal in the former case, intermediate for slower presses and maximal in the count-block and the look-only block.

The space bar was pressed on average about 250 ms after the onset of its illumination. Motor responses were thus output by the brain in 200 ms or less and thus before the onset of the N300 (Fig. 1). In accordance with the idea that the central N1 indexes an early automatic inhibition, the amplitudes of this ERP were found to be the largest for the two tasks where subjects had to prevent themselves from pressing the space bar. They were intermediate for the slow-press trials and smallest for the fast-press trials.

These effects appear remarkable because they were observed despite two factors moving ERPs in the direction opposite to the predictions. Firstly, pressing the space bar as fast as possible induced greater attention relative to the two other tasks and thus greater N1s, particularly at occipital scalp sites (Hillyard, & Anllo-Vento, 1998; Johannes et al. 1995; Hillyard et al. 1995; Luck, Fan & Hillyard, 1993, Luck, Heinze, Mangun & Hillyard, 1990), in accordance with the results observed in the present work at O1/2. Secondly, the space bar pressing done with the right hand induced motor potentials and thus greater negativities at the left central scalp sites (C3) just before the action and thus during the N1 time window (Coles, Gratton & Donchin, 1988; Miller & Hackley 1992). This is illustrated by Figure 1, where there was virtually no difference across the four conditions at this electrode site, and barely any at FC3, in the N1 deflections. At these sites, motor potentials have compensated weaker N1 processes. On the contrary, at right central sites, no or less motor potentials increased the negativity of ERPs of the N1 time window for fast press trials. Most likely, this is why much smaller N1 deflections were observed in this condition, in accordance with the early inhibition hypothesis.

The results thus strongly support the idea that the central N1s evoked by stimuli that are associated to actions indexes an early and automatic inhibition of the actions these stimuli just activated. The strength of this inhibition would fluctuate from trial to trial, allowing fast-presses when weaker, slower presses when stronger and preventing presses when maximum, as in the the count- and the look-only condition. On the other hand, the larger central N1s for the look-only- and for the count-task support the idea that the early and automatic inhibition is somewhat dependent on the task instructions and is boosted when subjects *have to* prevent the action. This task sensitivity would be consistent with findings showing that even automatic processes can be modulated to some extent by the task (e.g., Danion, 2013), just as the knee reflex can be prevented, reduced, or, on the contrary, amplified.

The second negative deflections, the N300s, appeared after the P3b, which can be the case in designs where only one stimulus is presented repeatedly and where no choice as to be made by participants (Donchin et al. 1978). N300s appeared in this study as large negative going deflections peaking around 400 ms post onset at central scalp sites. The lateness of their peak (i.e., 400 ms) could be due to the unusual presentations used here, that is, that of a stimulus suddenly occurring out of the dark, to which the subject was thus adapted during most of the experiment. These N300s were largest in the condition-blocks where presses had to be prevented, that is, for the look- and the count-task. This appears consistent with the idea that the N300 would, like the Nogo N2, indexes a late inhibition of actions, which is not automatic but occurs in tasks and contexts that require it and can relay the early inhibition indexed by the central N1. The N300 duration, which appears here to be longer than that of the N300s obtained for a unique stimulus repeated in an entire block of trials (Debruille et al. 2012), could be due to the prolonged duration of the presentations of the space bar. The brain continued to be activated by the presentation of a real 3D object for 500 ms, maintaining the need for inhibition.

Although a real object was used as a stimulus here, it was appearing in very special conditions. Only the spare bar was visible, not the rest of the keyboard. Moreover, the room was in complete darkness most of the times. Finally, the task of the subject was not to write a text and to press the space bar between two words, as is normally done with a space bar. It was to press it suddenly, completely outside of the context of its normal use. In such conditions, the appearance of the object may thus elicit an N300 even when the task instruction specifies to act upon the affordance and press the space bar as fast as possible. This could account for the remaining N300 deflections that can be seen in the ERPs of the press task and would be compatible with the idea that the N300 would, like the Nogo N2, indexes a late inhibition of actions, which is not automatic but occurs in tasks and contexts that require it. Note that, together with the attentional blink (Raymond, Shapiro & Arnell, 1992) and/or some motor refractory period, these remaining N300s activities might account for the absence of a second press in the case of fast-press trials. In effect, given that the space bar remained visible, it continued to activate the action of pressing it, which could have triggered a second press.

The large N1 and N300 differences found between fast- and slow-press trials reveal how much ERPs can differ from one trial to the next for the same stimulus presented in the same task. The inhibitory processes they index would thus spontaneously fluctuate in strength along time, just like vigilance. Another thing that may be inferred from the data is that even the late inhibition, that is, the one indexed by the N300s, would not last for a very long time. This can be deducted from the presence of the large N300s in the look-only- and in the count-condition, which show that N300s occur at many of the presentations of the space bar along the trials of these blocks. If the N300 inhibition had a long duration, no more N300 processes would have been needed after a few trials in these blocks and no large N300s would have been observed on the averages. The N1- and N300- inhibitions could provide the system with just enough time to fully process the meaning of the occurrence of the stimulus according to its context before acting. This meaning process is known to be indexed by the N400 ERP (for reviews see Kutas & Federmeier, 2011; Debruille, 2007 & Debruille et al. 2008; Shang & Debruille, 2013), which peaks around 450 ms for objects that are presented normally, that is, not in a complete darkness (see for instance Ganis & Kutas, 2003; Mudrik, Lamy & Deouell, 2009).

On the other hand, parietal P1s were marginally greater for fastest presses than for other condition-blocks. This is reminiscent of the larger centro-parietal P1 found by Kiefer et al. (2011). It thus seems possible to hypothesize that these centro-parietal P1s index the beginning of the activations of representations of the actions associated to stimuli. Further studies using stimuli activating actions less elementary than just pressing a computer key might be necessary to have larger P1 differences whose analysis will lead to clear-cut results. Note that the possibility that these greater P1s could, by overlap across participants, be responsible for the smaller central N1s can be eliminated by examining the sites at which no greater P1s can be seen for fast-presses and where N1s are still smaller (e.g., at FCz & FC4).

In conclusion, the results of this study suggest that components of the central N1 may index an immediate inhibition of actions whose prior activations could be reflected by greater parietocentral P1s. Further studies with methods such as the one used by Rabovsky and McRaey (2014) or by Cisek (2006), should be performed to see whether the amount of inhibition in the hidden layer(s) of their neuro-computational model could predict the size of the anterior N1s.

## Acknowledgements

This study was supported by the 2014-PR-171935 grant from the FQRNT allocated to J. B. Debruille, PI. We are thankful to Mathieu Brodeur and Paul Cisek for their inputs. We also thank Julie Henry for proof-reading the manuscript.

## Footnotes

1 Although Gibson (1977) first introduced the term to refer to the physical *properties* of the environment that are associated with actions, such as the grasping afforded by a stick or the warming afforded by a fire.

## References

Archibald, S. J., Mateer, C.A., Kerns, K. A. (2001) Utilization behavior: clinical manifestations and neurological mechanisms. Neuropsychology Review, 11(3), 117–130.

Atas, A. & Cleeremans, A. (2015). The temporal dynamic of automatic inhibition of irrelevant action. Journal of Experimental Psychology: Human Perception and Performance, 41(2), 289–305.

Bekker, E. M., Kenemans, J. L., Hoeksma, M. R., Talsma, D., & Verbaen, M. N. (2005) The pure electrophysiology of stopping. International Journal of Psychophysiology, 55(2), 191–198.

Binkofski, F., & Buxbaum, L. J. (2013). Two action systems in the human brain. Brain and Language, 127(2), 22–229.

Borghi, A., & Riggio L. (2008) Sentence comprehension and simulation of object temporary, canonical and stable affordances. Brain Research, 1253, 117–128.

Bruin, K. J., Wijers, A. A. (2002) Inhibition, response mode, and stimulus probability: A comparative event-related potential study. Clinical Neurophysiology, 113, 1172–1182.

Buxbaum, L. J., & Kalenine, S. (2010). Action knowledge, visuomotor activation, and embodiment in the two action systems. Annals of The New York Academy of Sciences, 1191, 201–218.

Chao, L. L. & Martin, A. (2000) Representation of Manipulable Man-Made Objects in the Dorsal Stream. Neuroimage, 12, 478–484.

Cisek, P. (2006) Integrated neural processing for defining potential actions and deciding between them: A computational model. Journal of Neuroscience, 26(38), 9761–9770

Clark, A. (1999) An Embodied Cognitive Science? Trends in Cognitive Sciences, 3(9), 345–351.

Coles, M. G. H., Gratton, G. & Donchin, E. (1988). Detecting early communication: Using measures of movement-related potentials to illuminate human information processing. Biological Psychology, 26, 69–89

Danion, F. (2013) Superposition of automatic and voluntary aspects of grip force control in humans during object manipulation. PLoS ONE, 11;8(11):e79341.

Debruille, J. B., Brodeur, M. B., & Franco Porras, C. (2012) N300 and social affordances: a study with a real person and a dummy as stimuli. PLoS One 7(10), e47922.

Debruille, J. B. (2007). The N400 potential could index a semantic inhibition. Brain Research Reviews, 56(2), 472–477.

Debruille, J. B., Ramirez, D., Wolf, Y., Schaefer, A., Nguyen, T. V., Bacon, B. A., … & Brodeur, M. (2008). Knowledge inhibition and N400: a within-and a between-subjects study with distractor words. Brain research, 1187, 167–183.

Donchin, E., Ritter, W., & McCallum, W. C. (1978). Cognitive psychophysiology: The endogenous components of the ERP. Event-related brain potentials in man, 349–411.

Electrode Position Nomenclature Committee (1991) American electroencephalographic society guidelines for standard electrode position nomenclature. Journal of Clinical Neurophysiology, 8, 200–202

Eimer, M., & Schlaghecken, F. (2003). Response facilitation and inhibition in subliminal priming. Biological Psychology, 64, 7–26.

Falkenstein, M., Hoormann, J. & Hohsbein, J. (1999) ERP components in Go/Nogo tasks and their relation to inhibition. Acta Psychol. (Amst), 101 (2–3), 267–291.

Franca M, Turella L, Canto R, Brunelli N, Allione L, et al. (2012) Corticospinal Facilitation during Observation of Graspable Objects: A Transcranial Magnetic Stimulation Study. PLoS ONE 7(11): e49025.

Ganis, G. & Kutas, M. (2003) An electrophysiological study of scene effects on object identification. Cognitive Brain Research, 16, 123–1444.

Gibson, J. (1977) The Theory of Affordances. In: Perceiving, Acting and Knowing: Toward an Ecological Psychology, Robert Shaw, John Bransford eds. Lawrence Erlbaum, NJ, p. 67–82.

Grafton, S. T., Fadiga, L., Arbib, M. A. & Rizzolatti, G. (1997) Premotor Cortex Activation During Observation and Naming of Familiar Tools. Neuroimage, 6, 231–236.

Greenhouse, G. W. & Geisser, S. (1959) On Methods of Analysis of Profile Data. Psychometrika, 24, 1582–1589.

Grèzes, J. & Decety, J. (2002) Does Visual Perception of Object Afford Action? Evidence from a Neuroimaging Study. Neuropsychologia, 40, 202–222.

Hillyard, S. A. & Anllo-Vento, L. (1998) Event-Related Brain Potentials in the Study of Visual Selective Attention. Procedures of the National Academy of Sciences, 95, 781–787

Hillyard, S. A., Mangun, G. R., Woldorff, M.G. & Luck, S.J. (1995) Neural Systems Mediating Attention. The Cognitive Neurosciences, 665–681.

Horváth, J. (2013) Action-sound coincidence-related attenuation of auditory ERPs is not modulated by affordance compatibility. Biological Psychology, 93(1), 81–87.

Jodo, E. & Kayama, Y. (1992) Relation of a negative ERP component to response inhibition in a Go/No-go task. Electroencephalography and Clinical Neurophysiology, 82, 477–482.

Johannes, S., Munte, T. F., Heinze, H. J., & Mangun, G. R. (1995) Luminance and Spatial Attention Effects on Early Visual Processing. Cognitive Brain Research, 2(3), 189–205.

Kiefer, M., Sim, E-J., Helbig, H. & Graf, M. (2011) Tracking the Time Course of Action Priming on Object Recognition: Evidence for Fast and Slow Influences of Action on Perception. Journal of Cognitive Neuroscience, 23(8), 1864–1874.

Kenemans, J. L. (2015). Specific proactive and generic reactive inhibition. Neuroscience & Biobehavioral Reviews, 56, 115–126.

Korb, S., Grangean, D., & Sherer, K. R. (2010). Timing and voluntary suppression of facial mimicry to smiling faces in a Go/NoGo task—An EMG study. Biological Psychology, 85, 347–349.

Kutas, M., & Federmeier, K. D. (2011). Thirty years and counting: finding meaning in the N400 component of the event-related brain potential (ERP). Annual review of psychology, 62, 621–647.

Luck, S., Fan, S. & Hillyard, S.A. (1993) Attention-Related Modulation of Sensory-Evoked Brain Activity in a Visual Search Task. Journal of Cognitive Neuroscience, 5(2), 188–195.

Luck, S.J., Heinze, H.J., Mangun, G.R. & Hillyard, S.A. (1990) Visual event-related potentials index focused attention within bilateral stimulus arrays. II. Functional dissociation of P1 and N1 components. Electroencephalography and Clinical Neurophysiology, 75, 528–542

Matheson, H., White, N., & McMullen, P. (2015) Accessing Embodied Object Representations from vision: A review. Psychological Bulletin, 141(3), 511–524.

Miller, J., & Hackley, S. A. (1992). Electrophysiological evidence for temporal overlap among contingent mental processes. Journal of Experimental Psychology: General, 121(2), 195.

Miller, E. K. (2000). The prefrontal cortex and cognitive control. Nature Reviews Neuroscience, 1, 59–65.

Mudrik, L., Lamy, D. & Deouell, L.Y. (2009) ERP evidence for context congruity effects during simultaneous object-scene processing. Neuropsychologia, 48(2), 507–517.

Niemi, P., & Näätänen, R. (1981). Foreperiod and simple reaction time. Psychological bulletin, 89(1), 133.

Ocampo, B., & Finkbeiner, M. (2013) The negative compatibility effect with relevant masks: a case for automatic motor inhibition. Frontiers in Psychology 4, 822.

Ocklenburg, S., Gunturkun O., & Beste, C. (2011) Lateralised neural mechanisms underlying the modulation of response inhibition processes. Neuroimage, 55 (4), 1771–1778.

Pires, L., Leitao, J., Guerrini, C., & Simoes, M. R. (2014) Event-related brain potentials in the study of inhibition: Cognitive control, source localization and age-related modulations. Neuropsychol Review, 24, 461–490.

Phillips, J. C. & Ward, R. (2002) S-R Correspondence Effects of Irrelevant Visual Affordance: Time Course and Specificity of Response Activation. Visual Cognition, 9(4-5), 540–558.

Proverbio, A. M., Del Zotto, M. & Zani, A. (2007) The emergence of semantic categorization in early visual processing: ERP indices of animal vs. artifact recognition. BMC Neuroscience, 8, 24, 1–16.

Rabovsky, M. & McRae, K. (2014) Simulating the N400 ERP component as semantic network error: insights from a feature-based connectionist model of word meaning. Cognition, 132(1), 68–89.

Raud, L., & Huster, R. J. (2017). The Temporal Dynamics of Response Inhibition and their Modulation by Cognitive Control. Brain Topography, 1–16.

Ridderinkhof, K. R. (2002a). Activation and suppression in conflict tasks: Empirical clarification through distributional analyses. In W. Prinz & B. Hommel (Eds.), Common mechanisms in perception and action: Attention & performance (Vol. XIX, pp. 494–519). New York, NY: Oxford University Press.

Ridderinkhof, K. R. (2002b). Micro- and macro-adjustments of task set: Activation and suppression in conflict tasks. Psychological Research, 66, 312–323.

Ridderinkhof, K. R., van den Wildenberg, W. P. M., Wijnen, J., Burle, B., & Posner, M. (Eds.). (2004). Response inhibition in conflict tasks is revealed in delta plots. Cognitive neuroscience of attention (pp. 369–377). New York, NY: Guilford Press.

Riddoch, M. J., Edwards, M. G., Humphreys, G. W., West, R. & Heafield, T. (1998) Visual Affordances Direct Action: Neuropsychological Evidence from Manual Interference. Cognitive Neuropsychology, 15(6-8), 645–683.

Rizzolatti, G., Luppino, G., & Matelli, M. (1998). The organization of the cortical motor system: new concepts. Electroencephalography and Clinical Neurophysiology, 106(4), 283–296.

Rizzolatti, G., & Matelli, M. (2003). Two different streams form the dorsal visual system: anatomy and functions. Experimental Brain Research, 153(2), 146–157.

Roche, R. A. P., Garavan, H., Foxe, J. J., & O’Mara, S. M. (2005) Individual differences discriminate event-related potentials but not performance during response inhibition. Experimental Brain Research, 160, 60–70.

Schneider, C., Lavoie, B. A., Barbeau, H., Capaday, C. (2004). Timing of cortical excitability changes during the reaction time of movements superimposed on tonic motor activity. Journal of Applied Physiology, 97, 2220–2227.

Schuck, S., Bayliss, A.P., Klein, C. & Tipper, S.P. (2010) Attention Modulates Motor System Activation During Observation: Evidence for Inhibitory Rebound. Experimental Brain Research, 205, 235–249.

Senderecka, M. (2016). Threatening visual stimuli influence response inhibition and error monitoring: An event-related potential study. Biological psychology, 113, 24–36.

Shang, M. & Debruille, J. B. (2013) N400 processes inhibit inappropriately activated representations: Adding a piece of evidence from a high-repetition design. Neuropsychologia, 51, 1989–1997.

Sumner, P., Nachev, P., Morris, P., Peters, A. M., Jackson, S. R., Kennard, C., & Husain, M. (2007). Human medial frontal cortex mediates unconscious inhibition of voluntary action. Neuron, 54, 697–711.

Symes, E., Ellis, R. & Tucker, M. (2006) Visual Object Affordances: Object Orientation. Acta Psychologica, 124, 238–255.

Raymond, J. E., Shapiro, K. L., & Arnell, K. M. (1992). Temporary suppression of visual processing in an RSVP task: An attentional blink?. Journal of experimental psychology: Human perception and performance, 18(3), 849.

Tipper, S.P. (2010) From Observation to Action Simulation: The Role of Attention, Eye-Gaze, Emotion and Body State. The Quarterly Journal of Experimental Psychology, 63(11), 2081–2105.

Tipper, S. P., Paul, M. A. & Hayes, A. E. (2006) Vision-for-Action: The Effects of Object Property Discrimination and Action State on Affordance Compatibility Effects. Psychonomic Bulletin & Review 13(3,) 493–398.

Tipper, S. P., Lortie, C. & Baylis, G. C. (1992) Selective Reaching: Evidence for Action-Centred Attention. Journal of Experimental Psychology: Human Perception and Performance, 18(4), 891–905

Tucker, M. & Ellis, R. (1998) On the Relations Between Seen Objects and Components of Potential Actions. Journal of Experimental Psychology: Human Perception & Performance, 24(3), 830–846

Vainio, L., Hammarén, L., Hausen, M., Rekolainen, E. & Riskilä, S. (2011) Motor Inhibition Associated with the Affordance of Briefly Displayed Objects. The Quarterly Journal of Experimental Psychology, 64(6), 1094–1110.

Wild-Wall, N., Falkenstein, M., Hohnsbein, J. (2008) Flanker interference in young and older participants as reflected in event-related potentials. Brain Research, 1211, 72–84.

Wilson, M. (2002) Six Views of Embodied Cognition. Psychonomic Bulletin & Review, 9(4), 625–636.

